# Evidence of hidden hunger in Darwin’s finches as a result of non-native species invasion of the Galapagoes cloud forest

**DOI:** 10.1101/2020.08.26.268136

**Authors:** Rebecca Hood-Nowotny, Ingrid Rabitsch, Arno Cimadom, Marcela Suarez-Rubio, Andrea Watzinger, Paul Schmidt Yáñez, Christian Schulze, Sophie Zechmeister-Boltenstern, Heinke Jäger, Sabine Tebbich

## Abstract

Invasive species pose a major threat to forest biodiversity, particularly on islands, such as the Galapagos. Here, invasive plants are threatening the remnants of the unique cloud forest and its iconic Darwin’s finches. We posit that food web disturbances caused by invasive *Rubus niveus* (blackberry), but also the management measures used to control it, could contribute to the rapid decline of the insectivourous warbler finch (*Certhidae olivacea*). We compared changes in long-term management, short-term management and unmanaged areas. We measured C:N ratios, δ^15^N-nitrogen and δ^13^C-carbon signatures in bird blood and arthropods, as indicators of resource use change, in addition to mass abundance and diversity of arthropods. We reconstructed the bird’s diets using isotope mixing models. The results revealed that finches in (*Rubus*-invaded) unmanaged areas foraged on abundant yet low quality arthropods and had shorter tarsi. Is this the first evidence of hidden hunger in degraded terrestrial ecosystems in Galapagos?

## Introduction

Invasive species are a major threat to biodiversity globally, even more to endemic island species which are particularly vulnerable, as host species gene pools and “escape” strategies are more restricted on insular island ecosystems (Atkinson, 1989). Species invasions at the primary producer level can cause massive ecosystem level changes (Szabo et al., 2012). In times of rapidly dwindling biodiversity, intensive habitat management is often the only option available to tackle invasive species and save the threatened focal species and/or their ecosystems (Moser et al., 2018). However, intensive management such as physical or chemical plant removal can also cause ecosystem disturbances and have detrimental effects on non-target species. Assessing the direct and peripheral effects of management measures is difficult and labour- and time-intensive. Here we present a novel stable isotope approach that detects and diagnoses ecosystem degradation, allowing for rapid response actions. Disturbance of food web structures and niche structure degradation are implicitly preserved in the isotopic signature of the focal species, as isotopic ratio of an organism is the result of all trophic pathways making up that individual, reflecting the trophic niche (Layman et al., 2012).

Darwin’s finches, endemic to the Galapagos Islands, have inspired some of the most important concepts in evolutionary biology (Watson et al., 2018). Although, all 17 species (Lamichhaney et al., 2015) have evaded extinction since Darwin’s first voyage, recently, populations have been decimated by habitat loss and the introduction of two aggressive invasive species, *Rubus niveus* (blackberry) and the parasitic fly *Philornis downsi* (Cimadom et al., 2019). Significant population declines of Darwin’s finches in the highlands of Santa Cruz were identified from 1998 to 2014 (Dvorak 2012). The insectivorous warbler finch (*Certhidea olivacea*) has suffered the most, with a decline of up to 50% in the forested and 75% in agricultural areas (Mauchamp and Atkinson, 2010). The primary habitat of the warbler finch is cloud forest that has experienced a 99% reduction in its area since the middle of the last century (Dvorak et al., 2012) due to human agricultural activities but also due to invasion of introduced plant species such as the invasive blackberry *Rubus niveus* (Rentería et al., 2012). The cloud forest is dominated by the endemic tree species *Scalesia pedunculata* (Rentería et al., 2012) with max. height of 12 m and a breast height diameter of up to 30 cm (Jäger, unpubl. data). *Scalesia* is a member of the daisy family *Asteraceae* and grows in dense stands in the humid zones of the major Galapagos islands. Although on a decadal scale, *Scalesia* canopy-cover is not affected by the *Rubus* invasion, at the invaded sites, understory plant composition is dramatically altered; with impenetrable dense thickets of *Rubus* with an above ground biomass of up to 10t ha^-1^ (Rentería et al., 2012). We hypothesised that areas invaded by *Rubus* act as a plentiful food resource for the primary consumers, mainly arthropods and that the subsequent consumption by the finches of the available abundant “low quality” primary consumers, leads to trophic disturbances with physiological consequences for the insectivorous birds.

The Galapagos National Park Directorate has pursued a policy of intensive *Rubus* removal, with machetes and subsequent herbicide control since 2003, to protect the remaining Scalesia forest. The invasive species management leads to the temporary removal of the understory in the controlled areas and a reduced availability of arthropods (Cimadom et al., 2019). In the immediate aftermath of control measures, significant reductions in the warbler finch’s breeding success have been observed (Cimadom et al., 2014) suggesting a major disruption of the warbler finch’s food web structure (Cimadom et al., 2019). We posit that both the invasion of *Rubus* and the management of *Rubus* cause major ecosystem level changes in resource structures, which have significant implications for our focal species, the warbler finch (*Certhidea olivacea*).

We sought to determine whether measuring the isotope signatures and stoiciometry of blood samples from the focal species could be used as a metric to indicate habitat degradation, thus quantitatively characterizing trophic structures. Laymann (Layman et al., 2007) suggested, δ^13^C–δ^15^N niche space is a representation of the total extent of trophic diversity within a food web. This is based on the premise that organisms, consumers and prey species reflect the consequences of changes in their environmental conditions and habitat structure, revealing shifts in their diet through the isotopic signatures in their blood (Fry, 2006). We predict that changes in prey type and availability in the *Scalesia* forest, as a consequence of *Rubus* invasion or invasive species management, should be captured in the isotopic signatures of the blood of the adult insectivorous finches (Inger and Bearhop, 2008; Wessels and Hahn, 2010) following the maxim “You are what you eat...plus a few per mil” (Boecklen et al., 2011a; Fry, 2006; Wessels and Hahn, 2010).

Contingent on the food supply and choice, consumers will feed on different proportions of particular dietary components (Inger et al., 2006). If these components have different isotopic signatures, their contribution can be easily detected with isotope based statistical mixing models (Wessels and Hahn, 2010). To reconstruct the diet, the mixing models use the consumers’ isotope signature, the isotopic signature and elemental percentages of the dietary components and account for the trophic fractionation factor (the “...plus a few per mil “). The trophic fractionation represents the net-value between the consumer and diet signatures considering metabolic and physiological processes within the consumer (DeNiro and Epstein, 1978, 1981) In contrast to traditional methods such as stomach content analysis, stable isotopes provide information, not only on digested food, but also on the assimilated components (Caut et al., 2009).

Stable isotope analysis of diverse metabolically active tissues allows tracking of temporal changes in diet and integrates values over extended periods. A drop of blood reflects the isotope signature within a time-frame of weeks, (Wolf et al., 2009a) whereas feathers and bones changes over months-years. Isotopic signatures provide accurate information about the diet of organisms, and can reveal whether diets change due to migration, weather conditions, habitat degradation, age, fasting, moulting, etc (Boecklen et al., 2011b; Cherel et al., 2005; Hobson, 1999; Hobson et al., 1993; Jackson et al., 2012).

Furthermore, using the stable isotope signature, it is possible to determine the trophic position of different organisms in the food web. In general, there is a slight discrimination in the isotopic components (C and N) in animals with respect to their diet (DeNiro and Epstein, 1981, 1978). Trophic fractionation of nitrogen is generally higher than that of carbon and is caused by the preferential metabolism of light nitrogen compounds (Podlesak and McWilliams, 2006). The δ^15^N Range (NR) distance between two species, with the most enriched and most depleted values at opposite ends of the food chain, yields a representation of vertical structure within a food web. The larger the range, the more trophic levels and a greater degree of trophic diversity is generally assumed. This premise was adopted herein, suggesting that the convex hull area plotted in a δ^15^N-δ^13^C bi-plot of dietary components represents trophic diversity and thus niche space (Layman et al., 2007). Importantly, elevated ^15^N values in biological tissues are indicative of starvation, as a result of nitrogen and/or protein recycling during starvation (Fry, 2006). Stoichiometric information, particularly C:N ratios of blood, also provides information on feed quality in terms of protein content. Dietary crude protein content generally determines growth rates, specifically in chicks (Márcia et al., 2016). Low dietary protein density can lead to hidden hunger; the supply of sufficient calories, but insufficient protein-nutrients such as nitrogen (Gödecke et al., 2018) or micro nutrients, which may lead to growth impairment and stunting (WHO, 1995).

We predicted that changes in prey availability should be apparent in the isotopic signatures of the warbler finch’s blood. Analysing the blood isotope signatures of the finch populations should allow us to trace the consequences of ecosystem level changes in dietary resource structure (Boecklen et al., 2011), caused by the invasion and control of *Rubus.*

In an experimental set-up, we investigated the effect of different *Rubus* management strategies in different areas: heavily *Rubus*-invaded with no control measures (NC), areas where *Rubus* has been recently removed and managed, since 2015 (RC) and areas with long-term-*Rubus-* removal management, since 2012 (LTM). Specifically, we asked: Does arthropod biomass differ between management areas in different foraging strata and over the breeding season? Even if the three study areas have similar overall arthropod productivity, the proportion of arthropod species which are suitable as prey could be lower. Thus, we asked. Does the predominant prey consumed differ between the invaded and managed areas and is there a difference in prey quality? To address these questions, we used traditional gravimetric and abundance analysis, in combination with stable isotope and stoichiometric signatures. Specifically, we measured whether there is an overlap of carbon and nitrogen isotopic signatures between available prey and the warbler finch’s blood (after accounting for trophic fractionation) to gain an understanding of feeding pathways and total niche space. In addition, we obtained information on the prey quality i.e. carbohydrate/fats versus protein (C:N).

We hypothesized that the near-complete removal of the forest understory leads to a decrease in the quantity of arthropod prey available, and the presence of *Rubus* leads to an increase in available low quality arthropod prey, resulting in a decrease in bird-body mass index (BBMI), weight and tarsus length, as indicators of finches’ overall condition.

## Results

### Arthropod biomass

Biomass was measured in two sampling rounds in 2015; one at the beginning of the breeding season in late January (round 1) and one in the middle of the breeding season in mid-April (round 2). The wet and warm season usually begins in early December and precipitation tails off, usually finishing by the end of May, with the drier and cooler weather dominating for the rest of the year. Overall, arthropod biomass (dry weight) was highest in the long-term management area (LTM) and lowest in the recently controlled area (RC). When comparing the arthropod biomass across forest strata or layers, the canopy had significantly higher arthropod biomass than the other layers, moss and understory, in all cases (F_(2,162)_=25.323 P<0.001), (Figure 1). In the middle of the breeding season (round 2, mid-April) arthropod biomass was significantly higher than at the beginning of the breeding season (round 1, late January), (F_(1,162)_=5.104, P=0.025). Overall, recently controlled areas (RC) in round 1 had the lowest arthropod biomass and long-term managed areas (LTM) in round 2 had the highest (Figure 2).

**Figure 1.**
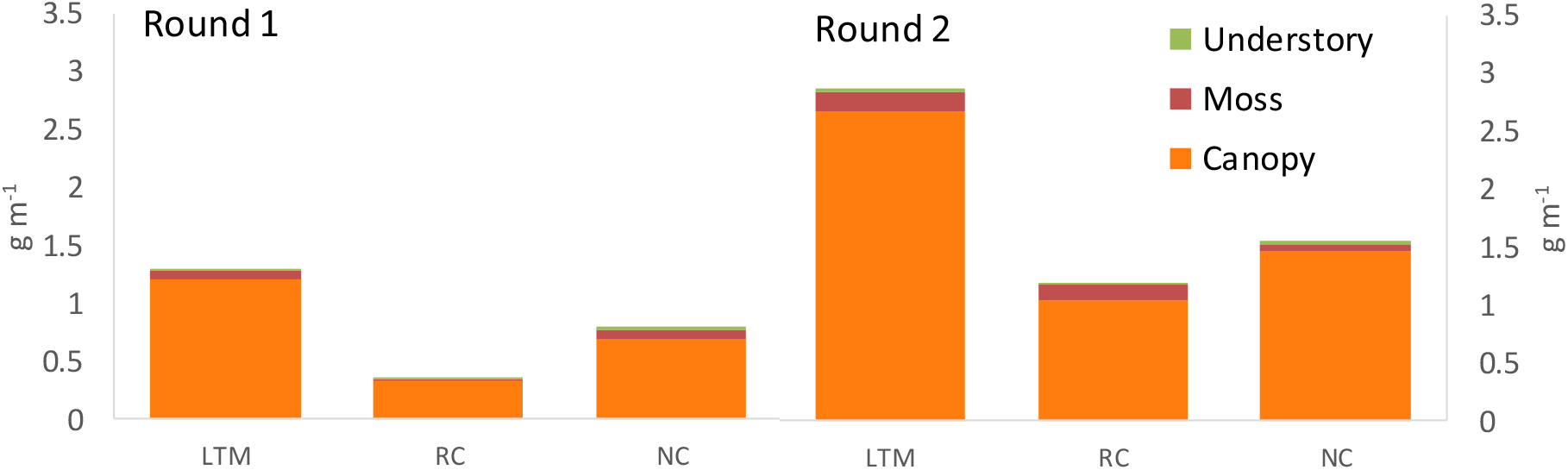
Total dry mass (g m^-2^) of arthropods per, management area, layer and round (n=10). Long-term management area (LTM), recently controlled area (RC) and unmanaged area (NC). Bars: orange-canopy, red-moss, green-understory. All values excluding Diplopoda, 2015.

**Figure 2.**
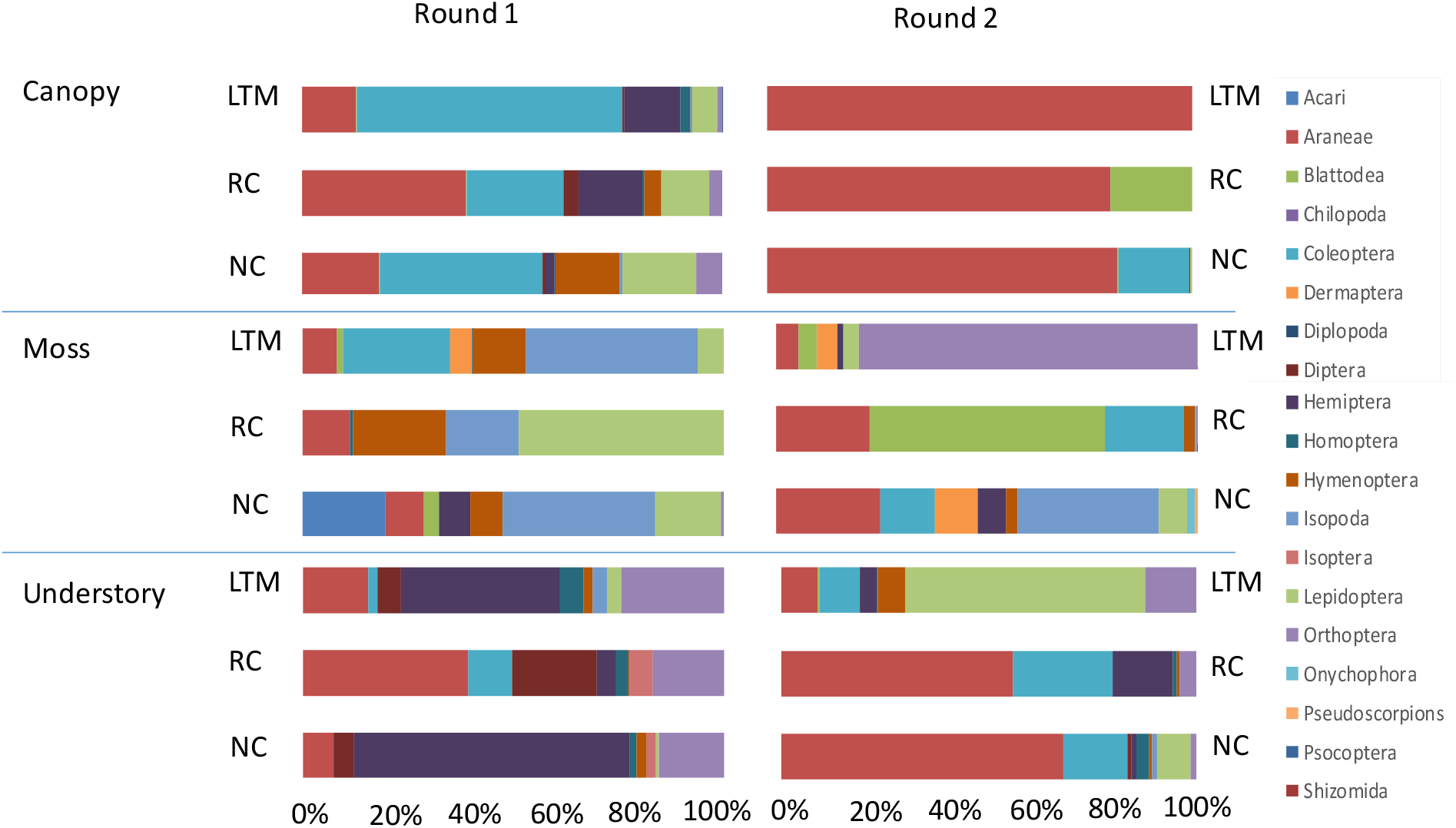
Relative dry mass abundance of arthropod orders per management area across rounds. Long-term management area (LTM), recently controlled area (RC) and unmanaged area (NC). All values excluding Diplopoda, n=10, 2015.

We discounted Diplopoda from our analysis, as observations had shown, from focal follows monitoring bird foraging, as well as bird stomach contents analysis, that due to their large size, Diplopoda were never eaten by the finches. Following convention, once it has been established that birds do not feed on a specific species, herein Diplopoda, it is reasonable that they can be excluded from the investigation (Wolda, 1990)

Relative arthropod biomass data showed similar relative abundance patterns within the forest layers across management areas. When comparing dominant arthropod orders between rounds and forest layers, we found that in each sampling round, each forest layer had consistently similar dominant arthropod orders (Figure 2). However, the dominant orders (ranked by dry weight biomass) differed between forest layers. In the canopy, the two dominant orders were Araneae and Coleoptera in all three study areas. In the two areas (RC & LTM), the third most important order was Hemiptera, which were almost absent in the unmanaged area (NC). In the unmanaged area, the third most important order was Lepidoptera.

In the moss layer, the dominant arthropod orders differed between the three study areas: In the recently controlled area (RC), the most dominant orders were Lepidoptera, followed by Hymenoptera and Araneae. In the long-term-managed area (LTM), dominant orders were Coleoptera and Hymenoptera followed by Araneae. In the un-managed (NC) the most dominant order was Acari, followed, by Lepidoptera and Araneae.

In the understory from the recently controlled area (RC), the dominant order was Araneae, followed by Diptera and Orthoptera. In the long-term-managed area (LTM), the dominant order was Hemiptera, followed by Orthoptera and Araneae. In the unmanaged area (NC), the order of the dominant orders was the same as in the long-term management area (LTM) but the relative mass abundance of Araneae was much lower than in the other two areas (less than 10%).

### Primary producer isotope signatures

*Scalesia pedunculata* was set as the isotopic dietary baseline. The δ^15^N isotopic signatures of both the *Scalesia* and the *Rubus* leaves were not significantly different across the three different management areas but nitrogen isotopes signatures of the two species were significantly different from one another (*Scalesia* mean: 5.7‰ and *Rubus* mean: 1.2‰, F_(1,57)_=139 p<0.001). There were significant differences in δ^13^C of *Scalesia* across sampling events of the different rounds, but not between management areas, in the drier-latter part of the breeding season-round 2 values were more enriched (F_(1,24)_=18.56 p<0.0001 Figure S2), this was attributable to differences in seasonal plant water availability and had no influence on the consequent dietary reconstructions. There was no significant difference between the molecular C:N ratios of *Rubus* and *Scalesia* leaves.

### Arthropod isotope signatures and quality

The arthropod weighted average δ^15^N’s were significantly different across management areas (F_(2,159)_=3.765, P=0.025) and forest layers (F_(2,159)_=14.098, P<0.001), but not across rounds (F_(1,159)_=0.648, P=0.422), (Figure 3, Table 1). Multiple comparison analysis highlighted significant differences in weighted average arthropod δ^15^N between the unmanaged area (NC) and the short-term management area (RC), (Tukey HSD-P_adj_=0.025).

**Figure 3.**
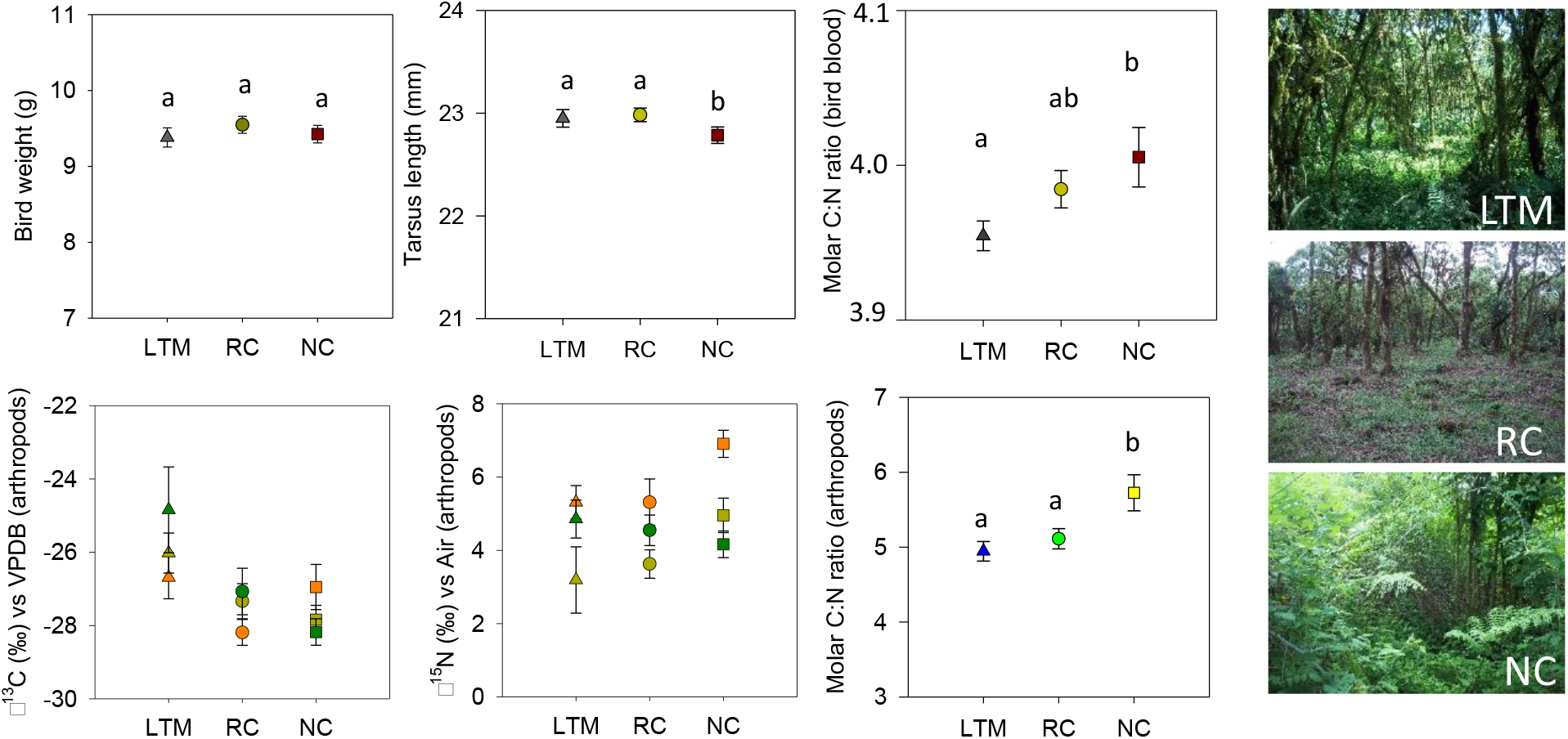
Upper panel: Birds’ mean weight (left), mean tarsus length (middle) and mean C:N ratio (right) of finch blood (in all cases mean±SE, 2015 & 2016, note multifactoral analysis suggested there was no significant differences between years so data were combined, n=60), letters indicative of Post Hoc-Tukey HSD Test. Lower panel: Arthropods’ mean weighted δ^13^C (left), mean weighted δ^15^N (middle) values excluding diplopoda and molar C:N ratio of arthropods (right, mean ±SE, 2015, both rounds, note multifactoral analysis suggested there was no significant differences between rounds so data were combined, n=60). Colours represent Forest Layers: canopy (orange), moss (dark yellow) and understory (dark green). Triangles: LTM, circle: RC and square: NC, means ±SE Right-hand panel photographs of management Areas: Long-term management area (LTM), recently controlled area (RC) and unmanaged area (NC).

**Table 1:** Isotope and elemental analysis of all arthropods’ δ^13^C, δ^15^N and C:N ratios (variables), ANOVA P-values. Factors: round, management area and forest layers.

| | $\delta^{13}\text{C}$ P-value | $\delta^{15}\text{N}$ P-value | C:N ratio |
| --- | --- | --- | --- |
| round | **0.018** | 0.422 | 0.620 |
| Area | 0.082 | **0.025** | **0.002** |
| Forest Layer | **0.004** | **<0.001** | **<0.001** |

Significant effects printed bold.

Arthropod’s C:N ratios were significantly different across management areas (F_(2,158)_=6.340, P=0.002), (Figure 3), and across forest layers (F_(2,158)_=9.152, P<0.001). Pairwise comparison showed that values of C:N ratios from the unmanaged area (NC) were significantly higher than values in the managed areas, RC and LTM (P=0.002 & P=0.023, respectively). Canopy C:N ratios were significantly lower than the other two forest layers, moss (P<0.001) and understory (P=0.047). Nitrogen densities (mg N m^-2^) were significantly different across forest layers (F_(2,158)_=20.280, P<0.001). Canopy arthropods had significantly higher nitrogen densities than the other two forest layers, moss and understory (P<0.001 in both cases). Although whole system mean arthropod nitrogen densities ranged between 3.97 and 8.31 mg N m^-2^ for the three management areas) significant differences were not detected, possibly a consequence of high variation and compounding measurement uncertainties.

The trophic structures of the *Scalesia* forest persisted across the management types based on the isotope signatures, warbler finches occupied the highest position in the trophic web sampled. As predicted, arthropods, in general, occupied the lower levels, the herbivorous arthropods consistently had the lowest δ^15^N values and the carnivorous arthropods “a few per mil” higher. Clear trophic isotopic enrichment was observed in the finch blood in the long-term management area (LTM), with the highest δ^15^N of and δ^15^N-range (of all the areas, figure 5) with less distinct differences between secondary consumers observed in the recently controlled area (RC) and unmanaged area (NC).

### Diet composition/ Diet selectivity?

No significant differences in δ^13^C of warbler finch blood were detected across management areas. However blood δ^13^C signatures were significantly different across years (F_(1,88)_=5.629, P=0.020); slightly more enriched in 2015 than 2016; rounds (F_(1,88)_=29.597, P<0.001), more enriched in round 2 than round 1; and sex (F_(2,88)_=8.800, P<0.001) less enriched in males than in females. Differences in signatures between rounds, within each year, were also significantly different in 2015 (P<0.001) and 2016 (P=0.013). The seasonal and annual differences are probably attributable to differences in plant water availability i.e. the effects of water stress on the plants cascading up through the arthropods, to the bird blood (Caut et al., 2009)

Multifactorial analysis revealed that bird blood δ^15^N signatures were significantly different across years (F_(1,88)_=5.861, P=0.018), 8.2‰ versus 8.6‰ in 2015 and 2016 respectively, rounds (F_(1,88)_=17.463, P<0.001), management areas (F_(2,88)_=18.107, P<0.001) and sex (F_(2,88)_=8.174, P<0.001) (Figure 5). Overall, values in the recently controlled area (RC) were significantly lower than those of the long-term management area (LTM) (P<0.001) and from the unmanaged area (NC) (P<0.001), as revealed by Tukey Post hoc (HSD) analysis (Figure 5). In 2015, only recently controlled area (RC) values were significantly higher (P<0.05) than in the long-term management area (LTM). In the unmanaged area (NC), only values from round 2 in 2016 were significantly different (P<0.001 for every case) from the other periods. Essentially, only in the unmanaged area (NC) did bird blood δ^15^N values change significantly between 2015 and 2016 (P<0.001).

Warbler finch dietary composition, as calculated based on the MixSIAR R-package, was different in each management area. In the recently controlled area (RC), dominant dietary components were Araneae, Hemiptera, Diptera and Lepidoptera, as predicted by the model (Table 2). The proportion of Diptera and Hemiptera was higher in the diet than expected from availabity, which shows that birds were clearly avoiding the Coleoptera. In the long-term management (LTM), Araneae and Hemiptera comprised over 70% of the diet, according to the model, and again, the warbler finch was not consuming the Coleoptera (Table 2). In the unmanaged area (NC), Hemiptera and Lepidoptera comprised 80% of the diet, based on the model, which was also a consiberably higher proportion of the diet than expected according to availabilty (Table 2) and again, also in this case appearing to select against the Coleoptera.

**Table 2.** Dominant dietary components (calculated using MixSIAR) of the warble finch compared with prey availability (arthropods relative dry mass abundance in forest) per management area based on round 1 data 2015.

| Orders | LTM |  | RC |  | NC |  |
| --- | --- | --- | --- | --- | --- | --- |
| | Diet proportion (mean ± SD %) | Availability (%) | Diet proportion (mean ± SD %) | Availability (%) | Diet proportion (mean ± SD %) | Availability (%) |
| Acari | 0±0.0 | 0 | 0±0.0 | 0 | 1±0.1 | 2 |
| Araneae | **41±0.4** | **12** | **34±0.4** | **38** | 5±0.2 | **17** |
| Coleoptera | 8±0.20 | **60** | 1±0.1 | **22** | 1±0.1 | **34** |
| Diptera | 1±0.1 | 0 | **19±0.3** | 4 | **11±0.3** | 1 |
| Hemiptera | **31±0.4** | **13** | **26±0.4** | **14** | **49±0.5** | 4 |
| Hymenoptera | 4±0.2 | 1 | 1±0.1 | 5 | 1±0.1 | **14** |
| Isopoda | 4±0.2 | 3 | 5±0.1 | 1 | 1±0.0 | 4 |
| Lepidoptera | 2±0.1 | 6 | **14±0.4** | **13** | **31±0.4** | **17** |
| Orthoptera | 9±0.3 | 1 | 0±0.0 | 3 | 0±0.0 | 6 |
|  | 100 | 97 | 100 | 99 | 100 | 99 |

High proportion values printed bold. Management areas: Long-term management area (LTM), Short-term management area (RC) and no-management (NC). Orange: Diet proportion. Green: Forest proportion.

Using the MixSIAR model analysis at the scale of primary producer, it was possible to determine dietary compositions from the individual forest layers, using the weighted average isotope values of the amassed collected arthropods as source inputs. Canopy arthropods were a dominant dietary source for the finches in the managed areas (LTM and RC), accounting for more than 95% of the warbler finch’s diet. On average, warbler finches were feeding almost exclusively from the canopy in the managed areas. However, in the unmanaged area (NC), canopy arthropods made up only 47% of the diet with 52% of the dietary arthropods coming from the understory (Table 3.), according to the isotopic modelling.

**Table 3.** Dominant dietary sources (forest layer) within each management area, dietary proportion (%) calculated using MixSIAR (Mean ± SD), rounds 1 and 2, 2015.

| Forest layer | LTM | RC | NC |
| --- | --- | --- | --- |
| Canopy | 99±0.05 | 97±0.09 | 47±0.49 |
| Moss | <1±0.04 | <1±0.01 | <1±0.02 |
| Understory | <1±0.03 | 2±0.09 | 52±0.49 |

### Overall condition of warbler finches

Warbler finch body weight (9-11g) was not significantly different across years (F_(1,84)_=1.128, P=0.291), rounds (F_(1,84)_=0.507,P=0.479) or management area (F_(2,84)_=0.719, P=0.490). Although the unmanaged area (NC) had the lowest overall mean values (Figure 3). Females were consistently heavier than males and the birds deemed “unknown” sex (F_(2,84)_=23.967, P<0.001). Ratios of males, females and unknowns caught and sampled were similar for all areas, with on average about four males for every female and unknown caught.

There were no significant differences in bird-BMI (BBMI) across years (F_(1,80)_=0.001, P=0.975), rounds (F_(1,80)_=0.012, P=0.914) or areas (F_(2,80)_=1.008, P=0.369). The overall BBMI values were approximately 18 kg/m^2^ and warbler finches in the unmanaged area (NC) had a slightly higher values. Female’s BBMI was significantly higher than that of males (F_(2,80)_=21.451, P<0.001).

There was no significant or predictive correlation between δ^15^N-bird-blood and BBMI or bird weight, suggesting no strong indication of starvation from the δ^15^N-signal interactions (data not shown). However, a few δ^15^N values of higher than 10‰, possibly outliers, were observed in males in both the unmanaged area (NC) and long-term management area (LTM), these were retained in the analysis.

Warbler finch’s tarsus length (21-24mm) was significantly different across management areas (F_(2,84)_=3.369, P=0.039) and sex (F_(2,84)_=3.198, P=0.046). Warbler finches had significantly shorter tarsi (Student’s t test, T_(11,3)_=2.38, P=0.018) in the unmanaged area (NC) than in the managed areas (RC and LTM), (Figure 3), and females having smaller tarsus length overall.

## Discussion

In this study, arthropod biomass and isotopic data, combined with differences in δ^15^N, but not δ^13^C signatures, of the warbler finch’s blood across management areas suggested changes in the underlying food web structure (Figure 4). As hypothesised, the higher mean arthropod biomass, lower C:N ratio and higher δ^15^N-δ^13^C range in the long term-managed area (LTM) suggested that these areas had recovered or semi recovered their trophic structure, compared to the recently controlled (RC) and unmanaged (NC) areas with compromised niche structures. Classical ecological theory suggests that a sympatric species in established ecosystems have minimal resource use overlap, a consequence of competitive exclusion (Gause, 1934) explaining the broader niche space in the long term-managed area (LTM). This is in line with previous studies in benthic systems demonstrating that invasive species occupy a tighter isotopic niche space than their native counterparts (Jackson et al., 2012).

**Figure 4.**
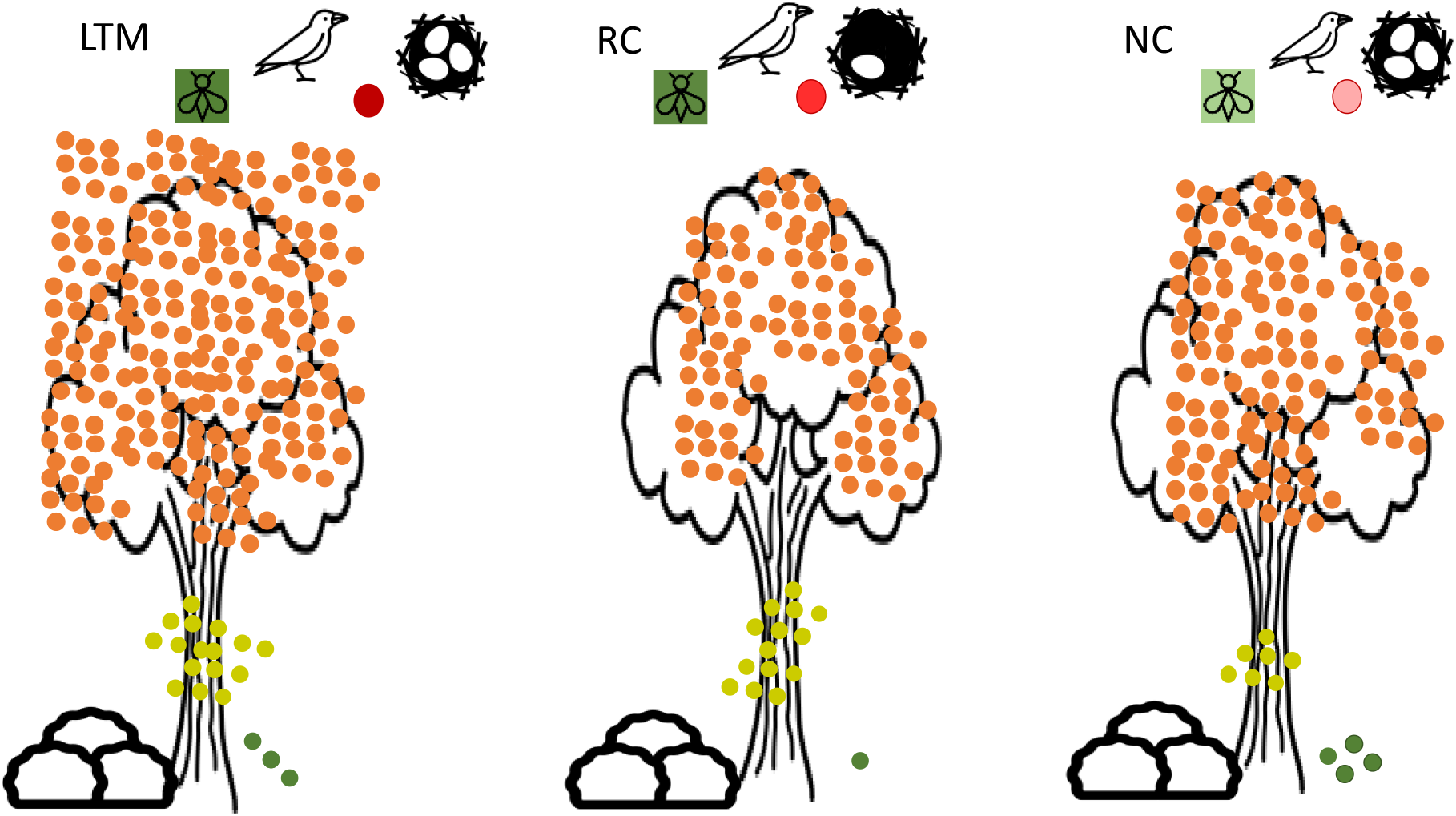
Schematic of whole system. Each spot in forest represents 10 mg dry mean mass of arthropods per m^2^, orange-canopy, dark yellow-moss, dark green-understory (nplots=10, round 2, 2015). Long-term management area (LTM), recently controlled area (RC) and unmanaged area (NC). Bug colour represents arthropod nutritional quality, with dark green high quality, light green lower quality. Red circles indicate bird blood stoichiometry, lower C:N ratio-high quality darker red, higher C:N ratio lower quality light red (nblood= 80). Eggs in nests indicate mean relative breeding success (Cimadom et al., 2019). Bird size highlighting lower tarsus length in NC area (but not to scale).

We scaled relative arthropod abundance measurements to absolute arthropod abundance per area measurements, based on dry weight data of the different arthropod orders per plot and layer. Although we did not capture all flying insects in our sampling strategy, we felt our methods enabled a reasonable estimate of the available prey biomass. Warbler finches are typical gleaners, not aerial hunters, which is consistent with observations and feeding frequencies in our study.

The isotope mixing model-MixSIAR analysis revealed that the bird’s dominant dietary components did not match the measured relative mass abundances of available arthropods in the long-term-managed area (LTM) and the unmanaged area (NC). This suggests a higher degree of prey selectivity in those areas and is consistent with the Optimal Diet Theory, which posits that predators chose prey that maximise their fitness (Christiansen et al., 1977; Endler, 1986). However, the available and consumed arthropod proportions were most similar in the recently-controlled area (RC). In addition, the lowest arthropod biomass was measured in the recently controlled area (RC), the highest arthropod biomass in the long-term managed area (LTM), indeed LTM biomass was three times that of the recently controlled areas (RC) and double that of the unmanaged areas (NC). These results suggest that low prey availability *per se* leads to less dietary choice, a phenomenon previously observed in long eared owls in Finland (Korpimäki, 1992). This comparison of isotope modelled-consumed versus available prey data is a useful metric, although rarely explored in the study of terrestrial ecosystems.

Arthropod biomass was ten times higher in the canopy compared to the moss and understory layers. It was significantly greater in round 2, than in round 1. This difference was most likely the result of the more humid conditions preceding the second sampling round, which is the typical climatic phenology of the Galapagos Islands (Petren et al., 1999, Grant, 1999, Grant et al., 2014). MixSIAR (Stock and Semmens, 2016) modelling revealed that in the managed areas LTM and RC, canopy arthropods accounted for more than 95% of the finches’ diet but in the unmanaged area (NC), it was only 41%, with a higher proportion coming from the understory (52%). This is corroborated by foraging observations that showed that warbler finches used the understory more frequently in the unmanaged areas (NC) than in the long-term-managed areas (LTM) (Filek et al. 2018). These results suggest habitat dependent food selection patterns and flexible feeding behaviours in these disturbed ecosystems. Similar shifts in change prey selection were observed in meso-carnivores in Chilean where forests were converted to plantations (Moreira-Arce et al., 2015). Our results also suggest that stable isotope signatures of focal species are a good indicator of niche disturbance.

The trophic consequence of the finches feeding predominantly on understory arthropods in the unmanaged area (NC) was detectable in the C:N ratio of the finch blood over both years and rounds. The C:N ratio of the blood is an indicator of diet quality (Gödecke et al., 2018) and was significantly higher in the unmanaged area (NC). These patterns were also observed overall in the C:N ratios of the arthropod samples, indicating that the arthropods, the warbler finches fed on, from in the unmanaged area (NC), were of significantly lower dietary quality, with higher C:N ratios lower and protein concentrations. There was no significant difference between the C:N ratios of *Rubus* and *Scalesia* leaves. This suggests that differences in arthropod C:N ratios, were a consequence of the lack of a secondary-arthropod-consumer in the trophic pyramid, the greater proportional abundance of available lower quality prey. A possible reason for this is that the trophic pyramid had not had sufficient adaptive time to exploit all the available niches in accordance with the classical ecological theory of competitive exclusion (Gause, 1934). In the unmanaged area (NC), the consistently lower δ^15^N range (Figure 5), specifically the tighter trophic distance between secondary consumer (δ^15^N carnivorous arthropod) and apex consumer in this system (δ^15^N_bird_), as well as the lower standard deviation of δ^15^N of the components therein, suggested a more constrained trophic structure, corroborating this finding (Layman et al., 2007).

**Figure 5.**
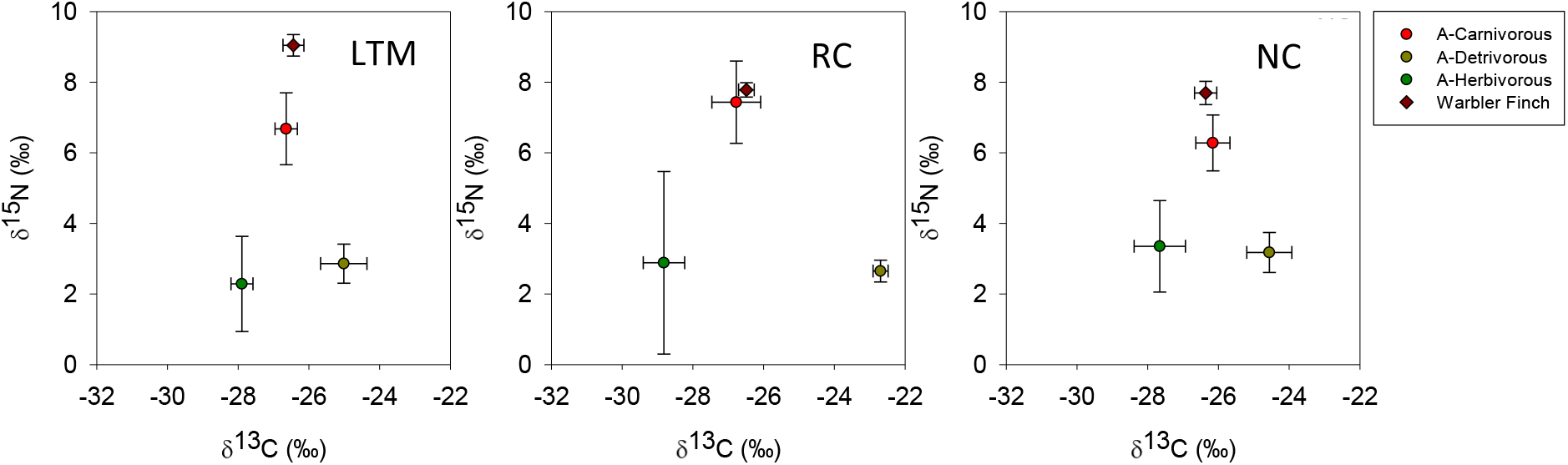
Scatter biplots of trophic web representation in the management areas. Isotopes signatures (“raw”-not accounting for trophic fractionation factor (TFF)), round 1, 2015. Management Areas: Long-term management area (LTM), recently controlled area (RC) and unmanaged area (NC). Dark red diamond: warbler finch. Red circle: carnivorous arthropods (Araneae). Green circle: herbivorous arthropods (Lepidoptera and Orthoptera). Brown circle: detrivorous arthropods (Diplopoda and Isopoda).

This dietary niche shift in the arthropod species as a result of the *Rubus* invasion indicates the collapse of the native food web structure, in which finches in the unmanaged area (NC) shift to feeding dominantly directly on primary consumers. Indeed, both the δ^15^N isotopic signatures of the understory arthropods and the reduction in the quantity of high quality Araneae, from the measured biomass and predicted dietary values from the MixSIAR analysis, of the finch diet in the unmanaged area (NC) suggest this. This is further substantiated by the lower mean nitrogen density values of the arthropods from the unmanaged area (NC). The MixSIAR analysis suggests that in the unmanaged area (NC), nearly half of the warbler finch diet consists of Hemiptera, which were abundant in the understory while Araneae were less abundant in the understory of the other two treatment areas. In many passerine birds, Araneae-spiders form an important and high-quality component of chick’s diet (Magrath et al. 2004), especially during early stages of chick development (Cowie and Hinsley, 2009; Grundel and Dahlsten, 1991; Naef-Daenzer et al., 2000). Spiders contain high level of taurine (Ramsay and Houston, 2003), which has multiple vital roles in the early development (Aerts et al., 2002) and is required for normal growth as well as the development of brain and visual systems.

Differences in finch diet quantity and quality led to significant differences in warbler finches’ size. Finches had significantly shorter tarsus length in the unmanaged area (NC) than in the managed areas (RC and LTM), but there were no significant differences between rounds or years (Figure 3). We argue that tarsus length is a more robust measure of long-term nutritional status than bird weight or BBMI, as it is an integrated measure and not subject to daily variations due to environmental or physiological status (Kempster et al., 2007). Evident from the fact that despite the differences in finch diet quantity and quality, there were no significant effects on bird weight or bird-BMI, between areas, years or rounds.

Warbler finches also had significantly lower breeding success in the short term management areas (RC), evidently a consequence of lower total arthropod mass (Cimadom et al., 2019). Finch breeding success was higher in both the unmanaged and long-term managed areas (Cimadom et al., 2019). This suggests that despite lower food quality in the unmanaged area, breeding was not affected. The shorter tarsus length, however, could indicate that parents compensated quality with quantity, fulfilling the chick’s calorific needs but not necessarily their nutritional requirements, at the expense of the size of the chicks. Low protein conditions and lack of nutrient funnelling may have caused the shift towards smaller finch size and shows parallels to human hidden hunger (Gödecke et al., 2018). An alternative explanation is that smaller finches were competitively driven out from the higher quality habitats. In the short term management area (RC), quantity of food rather than quality appeared to be the dominant constraint.

Taken together, our data herein suggest that there is a trophic pyramid collapse, due to the invasion of *Rubus*; a bottom-up control on ecosystem productivity and quality. This shift to a low quality diet was evident in both the isotopic and stoichiometric signatures of the warbler finch’s blood and arthropod biomass we posit that it subsequently influenced warbler finch’s size. We suggest that rapid environmental change due to the *Rubus* invasion did not allow for the finch population or the lower orders in the food web to adapt or adjust to the presence of the novel low quality diet.

Our date show that although management and control of invasive *Rubus* leads to dramatic temporary declines in food availability, these vital food resources can be re-established with persistent control measures and time. In this study, we demonstrate it is logistically feasible and financially possible to provide early warning signals of habitat degradation, using isotope and stoichiometric data, which can then provide management insights for effective ecosystem restoration.

## Materials and methods

### Study site

The study was conducted in the *Scalesia* forest at “Los Gemelos” (00°37’20” S, 90°23’00” W) on Santa Cruz Island, Galapagos, in 2015 and 2016. This area is dominated by the endemic tree *Scalesia pedunculata,* but most of its understory has been invaded by blackberry *Rubus niveus.* In some areas, the Galapagos National Park Directorate (GNPD) had controlled *Rubus* in order to restore the native forest. Within the forest, we defined three study areas, which differed in whether management of invasive *Rubus* took place and when: (1) the unmanaged area (NC, 8 ha), which was heavily invaded by *Rubus* and had never been exposed to any control measures; (2) the long-term manged LTM-area (6.7 ha), where *Rubus* had been manually and chemically controlled since 2012, and (3) the short-term controlled RC-area (6 ha), where *Rubus* had been controlled since August 2014 (for details see Cimadom et al. 2019). Control measures consisted of manually cutting *Rubus* with a machete and the subsequent application of herbicides (glyphosate and Combo©) on the regrowth (Schmidt Yáñez, 2016).

The wet and warm season characteristically starts at the beginning of December and ends by the end of May, the drier and cooler season persists for the remainder of the year (Jackson 1993). Different weather patterns were observed during the years 2015 and 2016. This was due to an El Niño event, which is an atypical wet season from September 2015 to the end of March 2016, followed by a dry season until the end of 2016 (supplementary figure S1).

### Study Design

There were four sampling campaigns in total, two rounds per year, which represented the start and end of the rainy season in typical years. round 1 was conducted from the end of January until the first week of February. round 2 was conducted in Mid-April.

**Figure S1.**
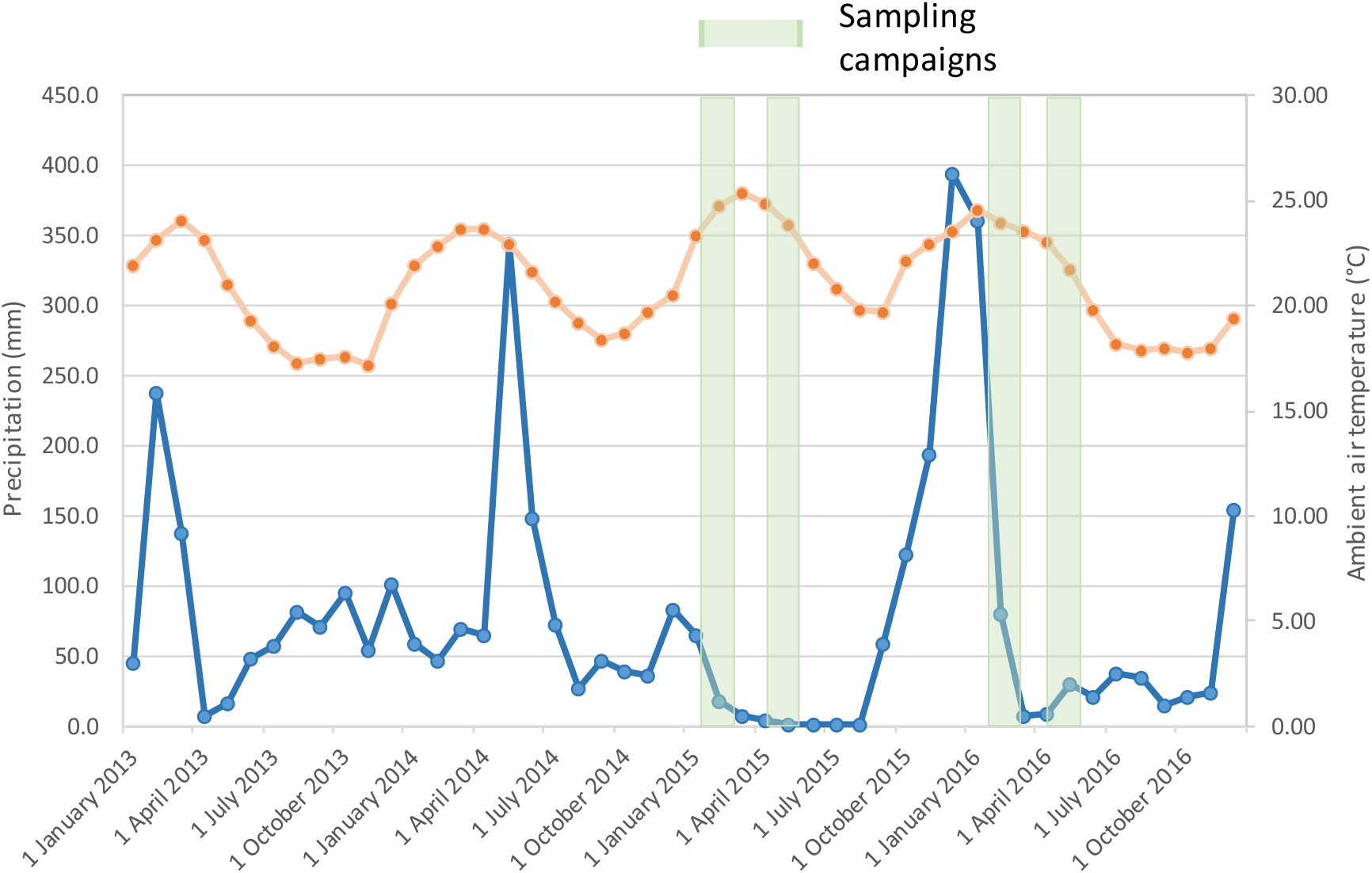
Sampling periods for finch blood, arthropod collection and monthly rainfall and temperature – from Los Gemelos, and El Carmen weather stations. The blue line represents monthly precipitation from 2013 until 2016. Dashed green columns represent the sampling events (round 1 and round 2) for 2015 and 2016. Orange line ambient air temperature, note 2015 months 01-09, temperature data are 30 year average values. Source: Charles Darwin Foundation.

Data on daily precipitation was provided by the nearest weather station (operated by Rolf Sievers, S 0°39’57.49’’ W 90°22’35.04’’) located about 4.5 km south and 150–200 m lower than the study site Los Gemelos. Precipitation data were available over the entire study period.

For each round in both years, **Warbler finch blood samples and arthropod samples** were collected from the three management areas. *Scalesia pedunculata* leaves from the canopy and *Rubus niveus* leaves from the understory were sampled randomly across management areas (5 replicates per area per round for *Scalesia,* 10 replicates for *Rubus*).

We collected ten replicate samples of warbler finch’s blood per management area (LTM, RM and NC), round (1 and 2) and Year (2015 and 2016), 120 samples in total. Blood samples were obtained by pinpricking the brachial vein of the finches with a lancet (Tebbich et al., 2004). One blood sample per individual was collected on a 5 mm^2^ Whatman GFA fibreglass filter discs, which was stored inside a coded test tube for subsequent isotope analysis.

### Birds were captured

with mist nets and ringed to avoid pseudoreplication. Left tarsus length was measured using a calliper (accuracy 0.01 mm). Each tarsus was measured twice, an average of the two values was used in our calculations. Birds were weighed using a field balance (accuracy 0.1 g). We developed a bird body mass index (B-BMI) based on the tarsus length and weight. We used an anologous formula to that of humans (WHO, 1995) weight (kg) /height (m)^2^, substituting height with tarsus length, as we used the tarsus length as a parameter for growth. All units were converted accordingly.

### We collected the arthropods

according to the collection procedure established by Schmidt-Yáñez (Schmidt Yáñez, 2016) from three defined microhabitats specifically canopy, moss and understory. Small tree finches and warbler finches mainly forage in the canopy, understory and in the moss growing on tree trunks (Filek et al. 2018) and we sampled arthropod biomass in each of these micro habitats. Canopy samples were taken by branch clipping. For this, a white polyester bag (diameter 50 cm, length 150 cm) was attached with clips to a metal ring (diameter 50 cm) at the top of a five-meter bamboo pole. The bag was pulled over a branch in the canopy (3-5 m height). The branch was then immediately cut off with a loppers (GARDENA), so that it fell into the collection bag, which was closed immediately by twisting the pole to prevent arthropods from escaping. Branches and leaves were then examined for arthropods inside the bag. All encountered arthropods were collected with an aspirator and stored in 70% alcohol. The branches and leaves were then put into a separate Ziploc bag for a second examination in the laboratory. The leaves were subsequently dried for 72 hours at ca. 60°C in a drying chamber to determine the dry weight.

Arthropods within the moss were collected from the same trees as the corresponding canopy samples. A 50 cm wide band of moss was carefully scratched off from the circumference of the tree trunk at a height of 1.5 m and transferred into a plastic tray. The moss was then briefly searched for larger arthropods that might escape from the tray area and then placed in a Ziploc bag for a second examination in the laboratory. As with the previous samples, all arthropods were stored in 70% alcohol. The moss samples were dried for 72 hours at ca. 60°C in a drying chamber to determine the dry weight.

To sample the understory, 5 m long transects with a buffer of 1 m width in each direction amounting to an area of 10 m^2^ were visually searched for 15 min by one person. Arthropods encountered on vegetation up to 1.7 m above the ground were collected either by hand or with an aspirator and stored in 70% alcohol. Flying insects could not be recorded by this method.

Standard methods to sample insects from understory vegetation (e.g. using a sweep net) could not be used, as the understory vegetation in our study area was invaded by spiny R. niveus. A canopy, understory and moss sample were collected in ten randomly selected sampling points in each of the three study areas We chose these microhabitats/forest layers because they were identified as the most important foraging substrates of the warbler finch (Filek et al., 2018). In 2015, we collected a total of 180 composite arthropod samples, ten replicates per forest layer (canopy, moss and understory), per management area (LTM, RC and NC), at two times (rounds 1 and 2) at the beginning of the breeding season late January (round 1) and in the middle of the breeding season in mid-April (round 2). The composite samples were created amassing the individual arthropods (all species), sampled at specific layer in a specific management area. All arthropods were collected regardless of their life stages and stored in 70% ethanol.

### Dry mass values of arthropods

were obtained by washing off the ethanol three times with deionised water and drying samples at 50°C overnight between each wash. This washing procedure had been tested and shown not to affect either δ^13^C and δ^15^N or nutrient content of the sample (Hood-Nowotny et al., 2016). Once dried, we weighed the samples (each arthropod order separately per field replicate) on a five-figure precision balance.

We identified samples from round 1 to arthropod order. We selected a representative group (individuals from one particular order) of each composite sample (from round 1) to be analysed for isotopic signature separately for further use in a diet reconstruction model. We chose the orders with the highest percentage of dry mass per sample, ensuring that there was at least one representative order (sub-sample) per forest layer or one per management area. Orders with less than 5% of dry mass out of the total mass of the composite sample, were not chosen for the individual isotopic analysis. With these representative orders, 162 additional sub-samples were created. We reunited the sub-samples with their respective analysed composite sample mathematically, by means of simple isotope based mass balance equations. This procedure was adopted to allow capturing the data in a logistically and economically feasible manner.

Once dry mass values were obtained, all composite samples and sub-samples were dried again, milled (Retch, DE) homogenised and a representative aliquot transferred (typically 3 mg) into 3.5 × 5 mm tin capsules, for analysis of stable isotopes of carbon and nitrogen, with a full range of standards bracketing all sample values. Subsequently, δ^13^C and δ^15^N of the sub-samples were back-calculated mathematically, using a simple mass balance equation and reunited individual sample to the corresponding composite sample to allow statistical analysis.

### IRMS samples were analysed

using a Flash 2000 Elemental Analyser in carbon and nitrogen configuration, linked to a Thermo Scientific Delta V Advantage automated isotope ratio mass spectrometer (IRMS) (Bremmen DE). A full complement of internal in-house and internationally certified standards was run with the samples to calculate isotopic ratios and % C and N values. The isotope ratios were expressed as parts per thousand per mil (‰) and as δ deviation from the internationally recognized standards Vienna Pee Dee Belemnite (VPDB) and AIR. All samples are referred to this scale from herein.

**Table S1.** Warbler finch’s blood and arthropods samples design (number of replicates)

| Sample type | Year | rounds | Forest Layer | Areas |  |  |
| --- | --- | --- | --- | --- | --- | --- |
|  |  |  |  | LTM | RC | NC |
| Warbler finch, blood,<br>Bird body mass index<br>(BBMI), tarsus length,<br>bird weight. | 2015 | 1 | - | 10 | 10 | 10 |
|  |  | 2 | - | 10 | 10 | 10 |
|  | 2016 | 1 | - | 10 | 10 | 10 |
|  |  | 2 | - | 10 | 10 | 10 |
| Arthropods | 2015 | 1 | Canopy (C) | 10 | 10 | 10 |
|  |  |  | Moss (M) | 10 | 10 | 10 |
|  |  |  | Understory (U) | 10 | 10 | 10 |
|  |  | 2 | Canopy (C) | 10 | 10 | 10 |
|  |  |  | Moss (M) | 10 | 10 | 10 |
|  |  |  | Understory (U) | 10 | 10 | 10 |
| *Scalesia* leaves | 2016 | 1 | Canopy (C) | 5 | 5 | 5 |
|  | 2017 | 2 | Canopy (C) | 5 | 5 | 5 |
| *Rubus niveus* leaves | 2017 | 1 | Understory (U) | 5 | 5 | 5 |

Management Areas: (LTM) Long-term management, (RC) Short-term management and (NC) unmanaged area.

**Figure S2.**
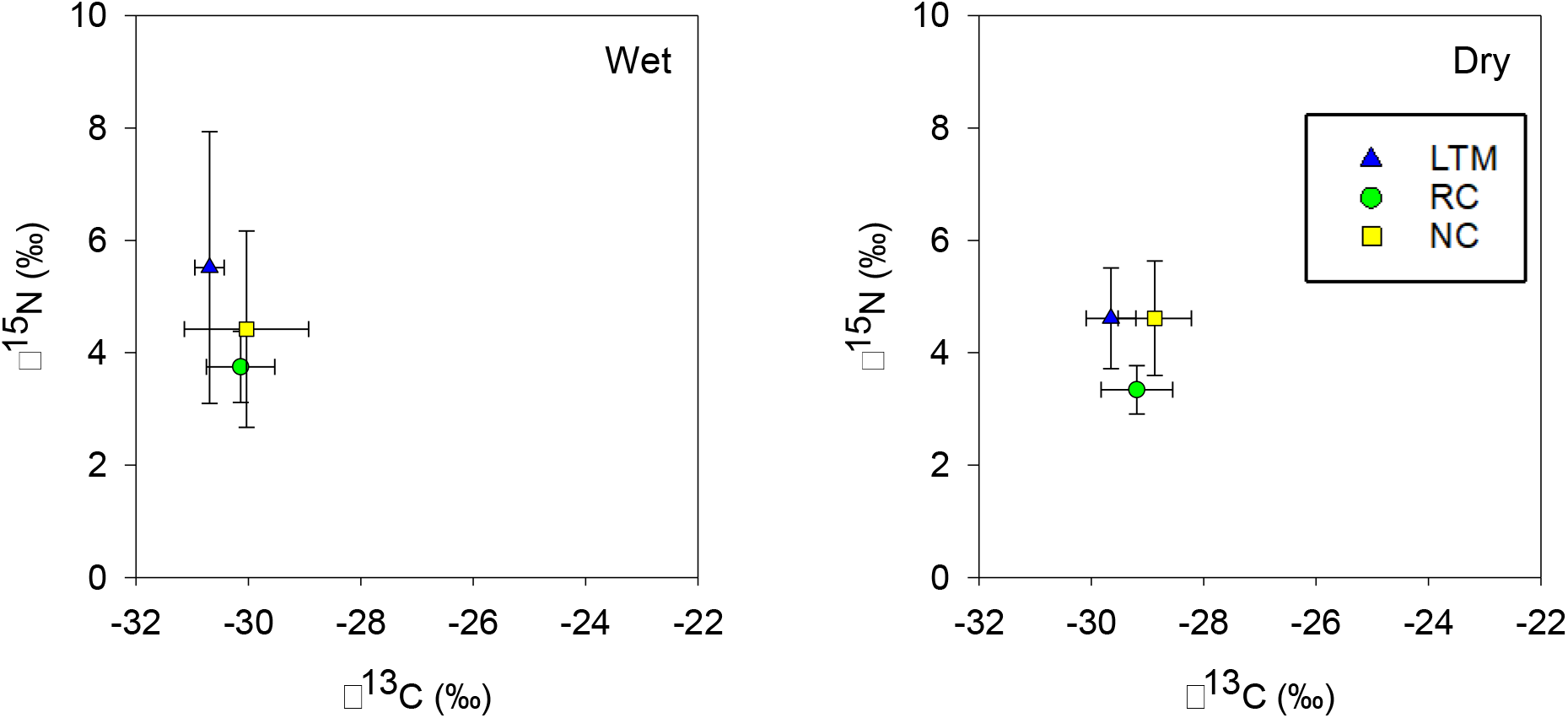
Scatter biplot of stable isotope signatures of *Scalesia pedunculata* (dominant tree in forest), **m**ean values (± SD) per management area (LTM, RC, and NC) across round. Management areas: (LTM) Long-term management, (RC) Short-term management and (NC) No management. Error bars represent the standard deviation.

### Arthropods Mass Abundance Standardisation

We standardised the arthropods mass abundance from the different forest layers to arthropod mass abundance per 10 m^2^ plot, which was the area of the sampled understorey plot. This was intended to achieve a better representation of the mass abundance of the available arthropods across the forest layers. Estimates of arthropod mass abundance were obtained for each forest layer, management area and round in the following way:

For the canopy arthropod samples, we applied a scaling factor according to Kitayama and Itow (Kitayama and Itow, 1999). Aboveground foliage biomass in a montane forest stand on Santa Cruz, Galapagos, was taken 1,482 kg of foliage per hectare, being 1,482 g of foliage in 10 m^2^. The correction factor related the arthropods mass and was scaled to the foliage mass measured of the leaves collected with the arthropods. The same leaf mass dependent scaling factor was used for all three areas since no significant differences were found in canopy cover between the areas in these experiments (Schmidt Yáñez, 2016).

For the moss arthropod samples, we applied a correction factor according to the surface area around the trunk, occupied by the moss collected. For this, we used the diameter at breast height (DBH) in meters measured for each sampling point. Knowing the surface (DBH × height) that a given mass of arthropods occupies at that sampling point, we estimated the arthropods mass per 10 m^2^. It was assumed that the percentage of moss cover did not change (between sampling points) and that the mass of moss varied proportionally. We did not apply a scaling factor to the understory data as the whole 10 m^2^ plot was sampled (Table S2).

**Table S2.** Correction formulas for estimating arthropod mass at each forest layer

| Forest Layer | Formulas |
| --- | --- |
| Canopy | $\text{Arthropod mass}_e = \frac{\text{Arthropod mass}_s}{\text{leaf mass}_s} * 1.482$ |
| Moss | $\text{Arthropod mass}_e = \text{Arthropod mass}_s * \left( \frac{10}{\pi * dbh * 0.5} \right)$ |
| Understory | No conversion made |

Note: (e) estimated value, (s) sampled value, all values were then divided by 10 to give values per square meter plot.

We excluded the order Diplopoda (generally, the millipedes) from the total mass data in 2015, as they have never been reported or observed to be consumed by the warbler finch (Filek et al., 2018). This assumption was supported by comparing the isotopic signatures of the Diplopoda with the finch blood data; the Diplopoda were well outside the sphere of consumption of the finches (Figure S3). Therefore, Diplopoda were excluded, since they sometimes dominated the samples in terms of mass (Table in Figure S3) and their presence was preventing us from teasing out the influence of the other orders.

**Figure S3.**
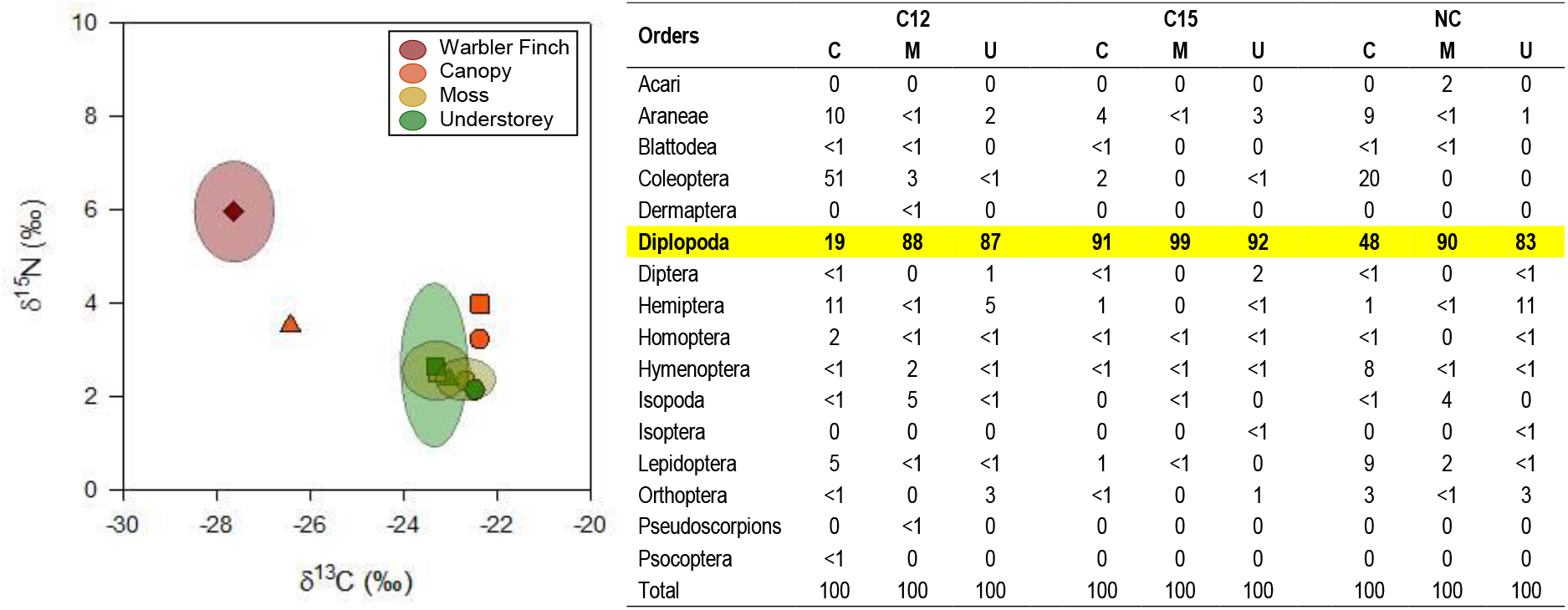
Scatter biplot of stable isotope signature of Diplopoda. Warbler finch represented by dark red diamond (overall mean value ±SD) **Diplopoda:** Shapes represent management areas: LTM: Long-term management area (triangle), RC: Short-term management area circle) and NC: unmanaged area (square). Colours represent forest layers. Orange: Canopy, yellow: Moss, green: Understory. Ellipses indicate standard deviation. Values in table represent percentage of total arthropod biomass (%). Table: Diplopoda is highlighted in bold and yellow. Forest Layers: (C) Canopy, (M) Moss and (U) Understory.

Exclusion of the Diplopoda from the data set allowed for a more nuanced analysis of the dietary data set. At the management area scale (amassing canopy, moss and understory samples), a ranking was made for each area type, roughly according to the Pareto (80:20) rule, leading to a top 3 dominant orders per area.

### Statistical Analysis

Multifactorial ANOVAs were performed on the following: the *Scalesia* samples, to evaluate the influence of rounds (1 and 2) and management areas (LTM, RC and NC) on δ^13^C and δ^15^N signatures; on the finch samples, to determine whether year (2015 and 2016), round (1 and 2), sex (male,female and unknown) and management area (LTM, RC and NC) had an influence on the bird blood data (δ^13^C, δ^15^N and C:N) and body metrics (weight, tarsus and bird BMI); on arthropods, to evaluate the influence of rounds (1 and 2), management areas (LTM, RC and NC) and forest layers (C, M, U) on each parameter δ^13^C, δ^15^N, C:N, nitrogen mass in 10m^2^ plot and dry mass. We used original isotopic signature values for the ANOVA analyses, i.e. signatures before subtracting values from the order Diplopoda to represent the isotopic structure of the *Scalesia* forest. Post-hoc analysis, Tukey HSD (Honestly-significant-difference) tests and Welch’s t-tests were performed, where applicable.

We compared both bi-plot C&N isotope signature data sets similar to Figure S3 (arthropods and bird blood), to evaluate the profile of the nutrients sources used by the birds and to determine whether the diet of the warbler finches reflected the arthropod signatures and which specific arthropod orders were dominant in their diet (data not shown). To determine potential diet components of the warbler finch, we analysed the isotopic signatures of all representative arthropods orders. Subsequently, we analysed management areas and forest layers for the most abundant orders and potential sources of food. We statistically analysed the generated signatures from the representative orders to determine if the orders had significant differences across management areas.

### Dominant Diet Sources and Components

We used a mass balance isotope mixing models to determine the composition of diets based on the dietary isotopic signatures. The warbler finch blood, representing the highest order consumer. The arthropods orders were defined as food sources with their corresponding trophic fractionation factors (Δ)(Hobson and Clark, 1992). We calculated diet proportion (*f*) and from isotopic values using the models.

To determine which of the arthropods orders present in the *Scalesia* forest were consumed by the warbler finch and in what proportion, we used Bayesian mixing models and compared the probabilities of all combinations predicting up to three possible dietary sources, under the creation of Markov Chain Monte Carlo (MCMC) chains. The analysis was conducted using the R-package “MixSIAR (Stock et al., 2016). MixSIAR creates and runs Bayesian mixing models to analyze biological tracer data (i.e. stable isotopes, fatty acids), which estimate the proportions of source (prey) contributions to a mixture (consumer). ‘MixSIAR’ is a framework that allows a user to create a mixing model based on their data structure and research questions, via options for fixed/ random effects, source data types, priors, and error terms (Stock et al., 2016).

To develop the MixSIAR model, we used the consumers’ signatures (warbler finch blood), possible food sources (orders signatures) and a trophic fractionation factor (TFF) for each element (carbon and nitrogen). The TFF used were obtained from the literature based on experimental values from laboratory studies on common quail’s blood (Hobson and Clark, 1992) as there were no equivalent values available for warbler finches. The models were created by establishing informative priors based on the relative abundance of the arthropods by taxonomic order and field observations (Filek et al., 2018). We set the factor managed area as a random effect, as we were analysing whether warbler finches fed on different components in different areas.

As diagnostic tools, we used the Gelman-Rubin Diagnostic and the Geweke Diagnostic. The Gelman-Rubin Diagnostic provides a value for each factorial. Less than 10% of those values should be below 1.05. The Geweke Diagnostic provides a standard z-score and 5% per chain and are expected to be outside +/-1.96.

We developed the set of food resources from the initial number of arthropod orders that were found in round 1 of 2015. Consequently, based on their relative abundance, we selected nine top orders, representing more than 95% of the total dry mass abundance (excluding Diplopoda). Several attempts were pursued to define priors based on abundance ranking and field observations, also increasing the length of the chain iterations. The best-fit model consited of more than 3000,000 chain iterations for the nine top orders.

We also used the MixSIAR package to determine which forest layer was the dominant diet source. We used the warbler finch blood data from 2015 (both rounds) for the consumer and we used the same TFFs. We entered the forest layers as sources. The best-fit model consisted of 1000,000 chain iterations for the three forest layers.

**Figure S4.**
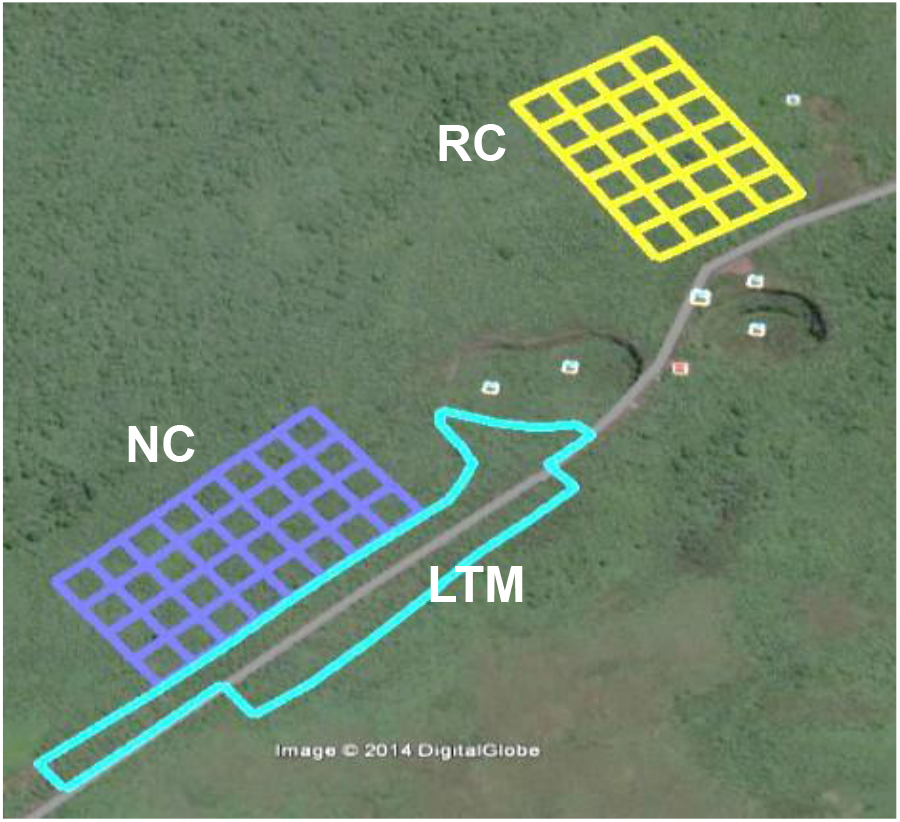
Site location. “Los Gemelos”, the road transecting the figure is the main road on Santa Cruz and distinct round structures are the extinct volcanoes.

**Figure S5.**
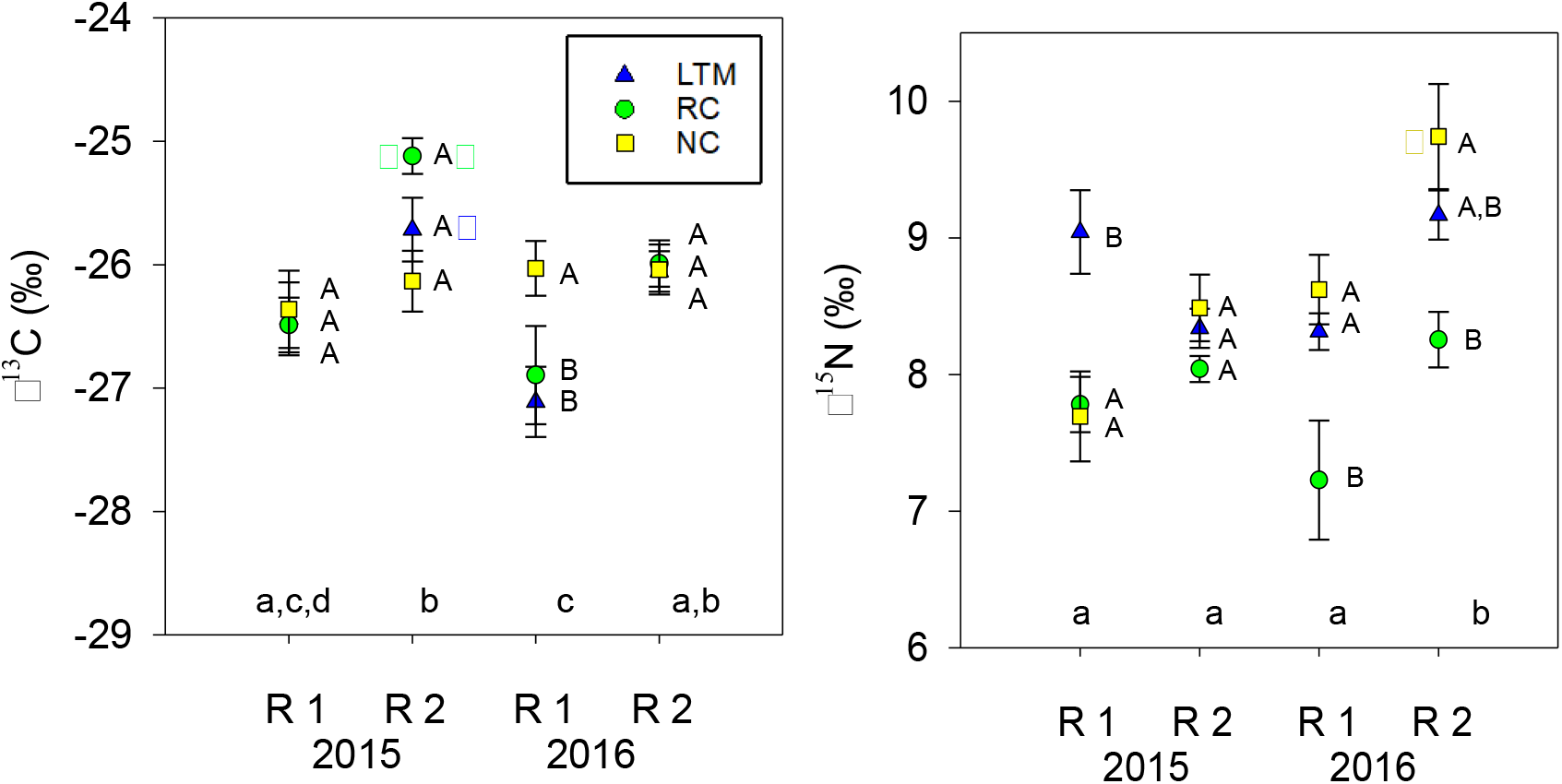
Stable isotope signatures of warbler finch blood. mean values (± SD) per management area (LTM, RC, and NC) across round. Management areas: (LTM) Long-term management, (RC) Short-term management and (NC) No management.

## Acknowledgements

This work was funded by the Austrian FWF P26556 and Galapagos Conservancy.

This publication is contribution number 2225 of the Charles Darwin Foundation for the Galapagos Islands.

## Author contributions

RHN & ST designed experiments. IR, AC, ST, CS, PSY, RHN, AW & HJ collected and analysed field samples and data. IR, RHN & MSR ran models and conducted statistical analysis.

RH and IR wrote first draft and received editorial input from ST, AC, MSR, SBZ & HJ. MSR & SBZ were the official supervisors of IR and RHN & ST the scientific supervisors.

